# Direct quantification of ligand-induced lipid and protein microdomains with distinctive signaling properties

**DOI:** 10.1101/2021.09.28.462247

**Authors:** Daniel Wirth, Michael D. Paul, Elena B. Pasquale, Kalina Hristova

## Abstract

Lipid rafts are known as highly ordered lipid domains that are enriched in saturated lipids such as the ganglioside GM1. While lipid rafts are believed to exist in cells and to serve as signaling platforms through their enrichment in signaling components, they have never been directly observed in the plasma membrane without treatments that artificially cluster GM1 into large lattices. Here we report that microscopic GM1-enriched domains can form, without lipid cross-linking, in the plasma membrane of live mammalian cells expressing the EphA2 receptor tyrosine kinase in response to its ligand ephrinA1-Fc. The GM1-enriched microdomains form concomitantly with EphA2-enriched microdomains, but only partially co-localize with them. To gain insight into how plasma membrane heterogeneity controls signaling, we quantify the degree of EphA2 segregation and study initial EphA2 signaling steps in both EphA2-enriched and EphA2-depleted domains. By measuring dissociation constants, we demonstrate that EphA2 oligomerization is the same in EphA2-enriched and -depleted domains. However, EphA2 interacts preferentially with its downstream effector SRC in EphA2-depleted domains. The ability to induce microscopic GM1-enriched domains in live cells using a ligand for a transmembrane receptor will give us unprecedented opportunities to study the biology of lipid rafts.

## Introduction

Lateral compartmentalization of the cell membrane into domains of different compositions and physical properties was first introduced to explain sorting of lipid and protein cargo to the apical plasma membrane in epithelial cells in 1988.^1–2^ Ten years later, the lipid raft hypothesis was formulated, proposing that domains enriched in sphingolipids, cholesterol, saturated lipids, and some membrane proteins form due to the phase behavior of lipids, and control key cellular processes like signal transduction, lipid and protein sorting, and viral entry during host cell infection.^3^ The lipid rafts have been further proposed to serve as platforms for signal transduction across the plasma membrane,^3^ bringing relevant receptors and downstream signaling molecules in close proximity.^4^ Studies in model systems like supported lipid bilayers, synthetic vesicles, and cell-derived vesicles have demonstrated that lipid heterogeneities can exist in membranes and can play a role in membrane organization and function.^5–12^ However, to this day the existence of lipid rafts remains controversial, as they have not been directly observed in unperturbed plasma membranes of live cells, without treatments that artificially cluster GM1 into large lattices using anticholera toxin anti-bodies.^10–12^

EphA2 is a member of the receptor tyrosine kinase (RTK) Eph family and has many important and diverse biological functions such as roles in blood vessel development, cell adhesion, and tissue patterning.^13–15^ The Eph receptors, including EphA2, are unique among the RTKs, because their biological ligands the ephrins are anchored to the surface of neighboring cells.^16^ Following contact with ligands such as ephrinA1 on neighboring cells, EphA2 has been reported to accumulate at contact regions between cells.^17^ In this study we show that we can induce microscopic domains that are enriched with the lipid raft marker GM1, as well as EphA2-enriched microdomains, by adding ephrinA1-Fc, a dimeric ligand which is known to emulate the function of surface-anchored ephrinA1 ligands in cell culture. We quantitatively characterize the domains in live cells and in model systems, we investigate the colocalization of GM1 and EphA2, and we probe key steps in EphA2 signal transduction across the plasma membrane, namely EphA2 association and adaptor protein binding. Furthermore, we test if EphA2 transmembrane (TM) domain sequence plays a role in the establishment of EphA2 heterogeneity in the plasma membrane. We report new observations and new insights into plasma membrane heterogeneities and their role in cell physiology.

## Results

### Formation of EphA2-enriched and GM1-enriched microscopic domains in the plasma membrane of live cells and vesicles in response to the ligand EphrinA1-Fc

Upon stimulation by surface-anchored ephrinA1, EphA2 has been reported to accumulate at contact regions between cells.^17^ The engineered soluble dimeric ligand ephrinA1-Fc (ephrinA1 fused to an antibody Fc region) is known to mimic the biological actions of ephrinA1.^18^ Here we first investigated if the addition of ephrinA1-Fc to cells expressing EphA2-mTurquoise results in accumulation of EphA2 in discrete regions of the plasma membrane. Experiments, shown in **Figure 1 a, i**, revealed that the ligand induces separation of EphA2 into EphA2-enriched and EphA2-depleted regions of the plasma membrane. We observed this behavior in both CHO and HEK293T cells. Quantification of EphA2 concentrations through measurements of mTurquoise fluorescence revealed that on average, the concentration of EphA2 in the EphA2 enriched regions is 2.4 ± 0.1 (range: 1.1 – 4.8) times higher than the concentration in the EphA2-depleted regions in HEK293T cells (**Figure 2 a**). Furthermore, **Figure 2 a** shows that the concentration of EphA2 in EphA2 enriched regions is 2.8 ± 0.1 (range: 1.0 – 6.4) times higher in CHO cells.

**Figure 1.**
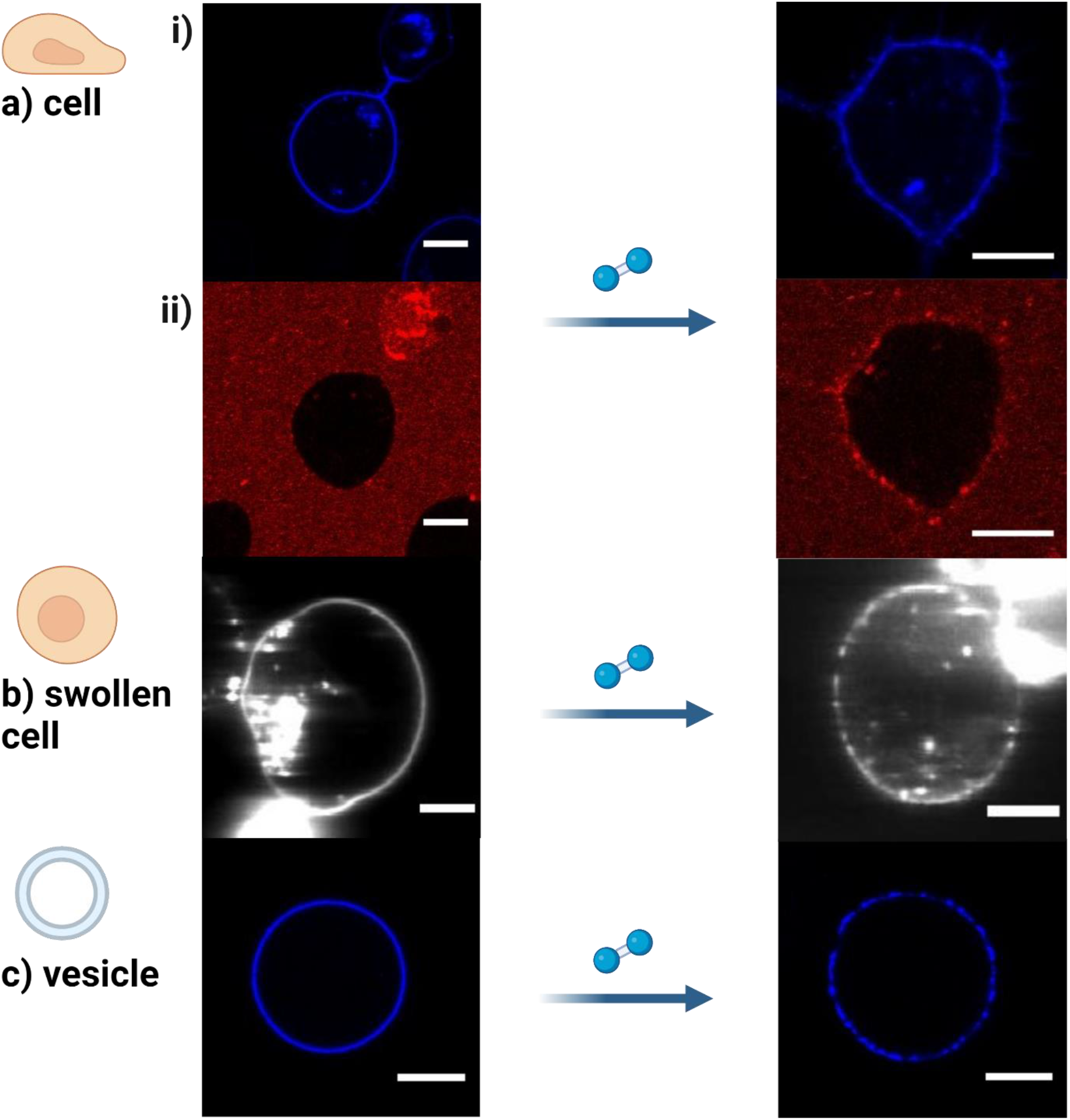
Induction of EphA2- and GM1-enriched microdomains. Addition of 50 nM ephrin-A1-Fc induces the formation of EphA2-enriched regions in a i) cells, b) swollen cells, and c) vesicles. a ii) shows the formation of GM1-enriched domains in cells after the addition of 50 nM ephrin-A1-Fc. The scale bar represents 10 μm.

**Figure 2.**
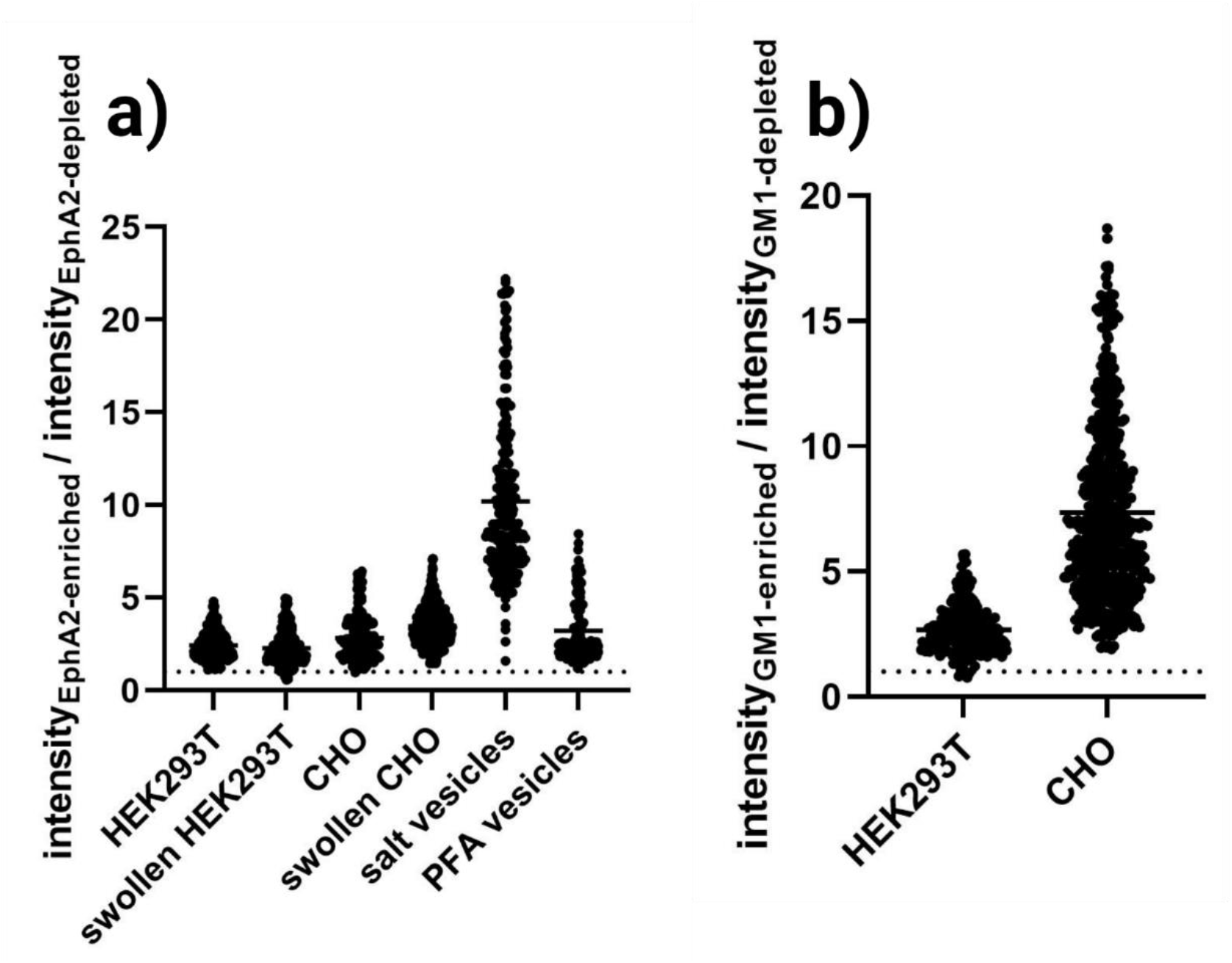
Fluorescence intensity ratios between enriched and depleted regions. Each ratio is calculated from enriched and depleted regions in the same cell. a) Intensity ratios of EphA2-mTurquoise fluorescence in HEK293T, swollen HEK293T, CHO, swollen CHO cells, osmotically-derived salt vesicles, and formaldehyde/DTT (PFA) vesicles. Notably, the intensity ratio for the osmotically derived vesicles is much higher than for cells. b) intensity ratios of CTB-AlexaFluor 555 fluorescence in HEK293T and CHO cells. The dotted line indicates y=1.

Next, we asked whether the EphA2-enriched regions might co-localize with GM1-enriched regions, a common marker for lipid rafts.^19–21^ We therefore added Alexa Fluor 555-labeled cholera toxin subunit B (CTB) to the EphA2 expressing cells, as CTB binds specifically to GM1.^22^ Experiments, shown in **Figure 1 a, ii**, revealed that the ligand induces separation of the membrane into GM1-enriched and GM1-depleted regions; the average ratio of CTB-Alexa Fluor 555 fluorescence in the GM1-enriched and depleted domains is 2.7 ± 0.1 (range: 0.8 – 5.7) for HEK293T cells and 7.4 ± 0.2 (range: 1.9 – 18.7) for CHO cells (**Figure 2 b**). To quantify the degree of co-localization of EphA2 and GM1, we performed two scans in a confocal microscope. In one scan, the mTurquoise bound to EphA2 was maximally excited, while in the second scan Alexa Fluor 555 on CTB was excited. No fluorescence bleedthrough between the two scans was observed. To quantify colocalization of EphA2 with GM1, Pearson’s correlation coefficient (PCC) was calculated using equation 1. A PCC of 1 indicates perfect positive correlation, a PCC of −1 shows perfect negative correlation, and a PCC of 0 shows no correlation at all.^23^ HEK293T cells and CHO cells were analyzed revealing PCC coefficients of 0.34 ± 0.04 (n=21) and 0.33 ± 0.02 (n=19), respectively.

A way to quantify GM1 affinity to EphA2-enriched domains (which is different than colocalization) is the so-called partitioning coefficient K_p,GM1_.^24–26^ To determine K_p,GM1_, the fluorescent intensity of CTB-Alexa Fluor 555 is quantified in EphA2-enriched and -depleted domains and divided to yield K_p,GM1_ (equation 2). Note, that K_p,GM1_ is different from the CTB-Alexa Fluor 555 fluorescence intensity ratios calculated above, since the regions are not directly chosen on the ‘GM1’-scan. Intensities used for the calculation were obtained in two different ways, as described in Materials and Methods: 1) by calculating means of annotated regions of interest (ROIs) (**Figure 3 a, ii**), or 2) by calculating intensity maxima along the intensity profile of selected lines (**Figure 3 b, i**). The results for the two different analysis methods are shown in **Figure 3 a and b, iii**, respectively. For both CHO and HEK293T cells K_p,GM1_ is > 1. Both analysis methods yield the same result for CHO (method 1: 1.7 ± 0.1; method 2: 2.0 ± 0.1) and HEK293T (method 1: 1.6 ± 0.1; method 2: 1.7 ± 0.1) cells. When compared to cells without added ephrinA1-Fc as controls (method 1: 1.01 ± 0.01 (HEK293T), 1.00 ± 0.02 (CHO); method 2: 1.04 ± 0.03 (HEK293T), 1.02 ± 0.04 (CHO)), the calculated K_p,GM1_ values are highly statistical significant (p_HEK293T_ < 0.0001, p_CHO_ < 0.0001 for **Figure 3 a and b**). This clearly shows that GM1 preferentially partitions into EphA2-enriched regions, as indicated by the colocalization analysis.

**Figure 3.**
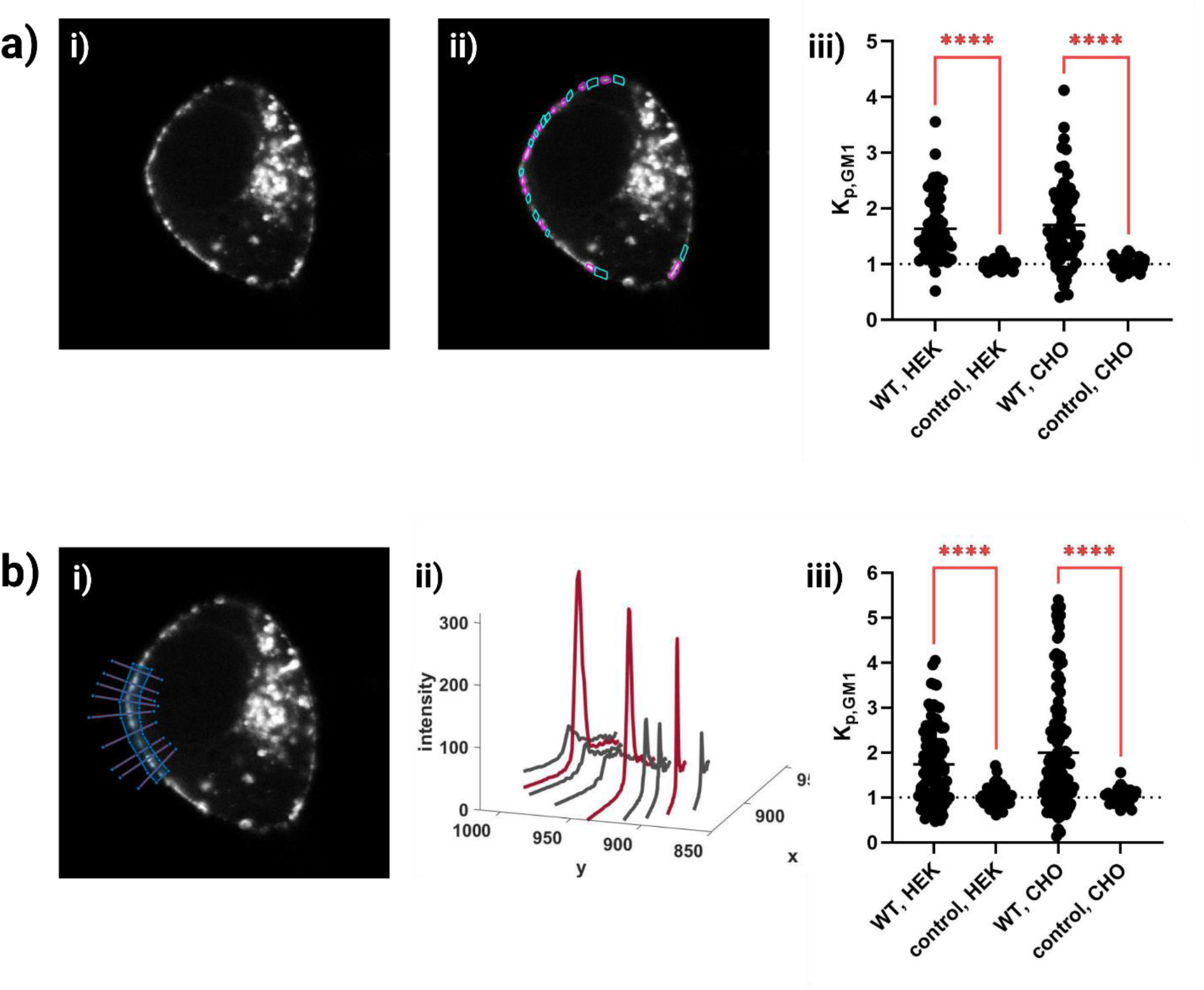
Calculation of the partition coefficient K_p,GM1_. Two different methods are used to calculate K_p,GM1_ according to equation 2. a ii) EphA2-enriched (pink) and -depleted (turquoise) regions are selected on the ‘EphA2’-scan. The corresponding mean CTB-Alexa Fluor 555 intensities are then used to calculate K_p,GM1_. b i) Lines that intersect with EphA2-enriched and -depleted regions are selected on the ‘EphA2’-scan. The maxima of their intensity profiles (b ii)) are used to calculate K_p,GM1_. The results for HEK293T and CHO cells are shown in a iii) and b iii). Cells without added ephrinA1-Fc serve as controls. Each K_p,GM1_ value comes from adjacent EphA2-enriched and -depleted regions within the same cell. The dotted line indicates y=1.

Next, we asked if the accumulation of EphA2 in discrete areas of the plasma membrane may be associated with endocytic processes. Typically, ligand-activated RTKs are uptaken into endosomes via a clathrin mediated mechanism, and thus it is possible that the EphA2-enriched regions we observe could be intermediates in the uptake pathway.^27–28^ As osmotic stress is known to eliminate endocytosis, we performed experiments in cells under reversible osmotic stress.^29^ Notably, the application of reversible osmotic stress is not lethal and completely reversible.^30^ Upon addition of ephrinA1-Fc, we again observed that the plasma membrane in cells under reversible osmotic stress separates into EphA2-enriched or depleted regions (ratio CHO cells = 3.47 ± 0.03, range: 1.5 – 7.1, **Figure 2 a**; ratio HEK293T cells = 2.3 ± 0.03, range: 0.5 – 5.0, **Figure 2 a**). As these cells have disrupted cortical actin (**Figure S1**), the observation in **Figure 1 b** suggests that neither endocytosis nor actin is driving the EphA2 separation. To further exclude actin and other cytoplasmic proteins as factors that contribute to EphA2 separation in the plasma membrane, we performed experiments with osmotically derived plasma membrane vesicles. These are ghost vesicles derived from CHO cells with no cytoskeleton and no cytoplasmic content.^31–32^ Indeed, cytoplasmic proteins, and proteins that are physically associated with the cytoplasmic leaflet of the membrane have been shown to leak out of these vesicles.^32^ Again, we observed that EphA2 in the vesicles separates into EphA2-enriched and EphA2-depleted regions (**Figure 1 c)**. The intensity ratio of EphA2-enriched and EphA2 depleted regions is 10.2 ± 0.3 (range: 1.6 – 22.2) (**Figure 2 a**).

Most of the studies investigating lipid rafts have been performed with plasma membrane derived vesicles, produced via DTT and formaldehyde.^33^ We asked if we will also observe high EphA2-mTurquoise intensity ratios in formaldehyde/DTT CHO vesicles. Surprisingly, 70% of the formaldehyde vesicles did NOT exhibit any EphA2 segregation. For the subset of formaldehyde vesicles which phase separated, we calculated intensity ratios of EphA2-enriched and EphA2 depleted regions with a mean of 3.2 ± 0.2 (range: 1.2 – 8.4) (**Figure 2 a**).

### The effect of lateral EphA2 segregation on EphA2 signaling

Next, we asked whether EphA2 signaling is different in EphA2-enriched and depleted regions. To investigate this, we probed lateral EphA2 self-association, as well as EphA2 interactions with an adaptor protein.

First, we asked whether the observed EphA2-enriched regions contain much larger oligomers than the EphA2-depleted regions. We thus measured EphA2 oligomer size in EphA2-enriched and depleted regions using Number and Brightness (N&B), which measures fluorescence fluctuations and reports on the oligomer size of a protein complex^34^. Experiments were performed in CHO cells under reversible osmotic stress, as described in^35^, where we imaged the equatorial plane of the membrane. Regions of EphA2-enrichment and EphA2-depletion were analyzed separately, yielding the oligomer size in these regions. **Figure 4** compares the oligomer size for LAT (a monomer control) with the EphA2 oligomer size in EphA2-enriched and -depleted regions. First, we see that EphA2 oligomer sizes measured in both EphA2-enriched (4.8 ± 1.1) and -depleted (3.4 ± 0.8) regions are significantly different when compared to LAT (0.9 ± 0.1), p < 0.0001 and p = 0.001, respectively. Second, we see much more significant heterogeneity in EphA2 oligomer sizes both in the EphA2-enriched and -depleted regions. Remarkably, we see no difference in the oligomer size between the EphA2-enriched and -depleted regions. The means of the oligomer size are 4.8 ± 1.1 and 3.4 ± 0.79, but the difference is not significant (p = 0.35). Thus, the oligomer size of EphA2 is similar in EphA2-enriched and -depleted regions and is characterized by a high degree of heterogeneity in both regions.

**Figure 4.**
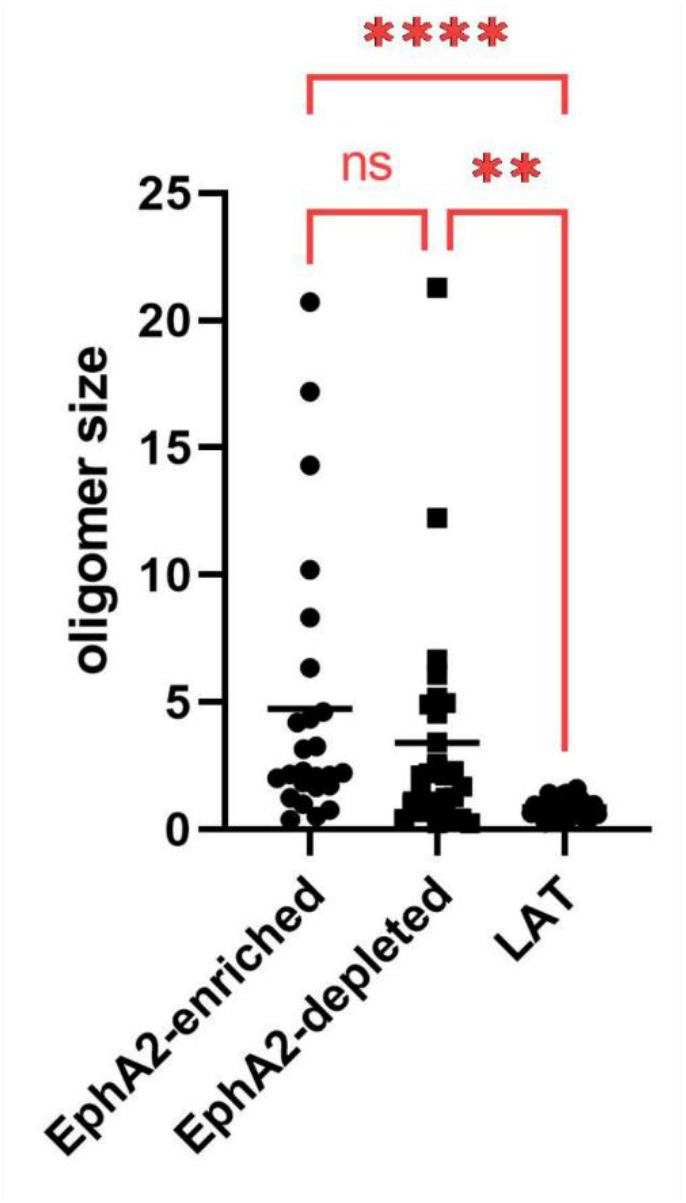
Number and Brightness (N&B) measurements of EphA2 oligomer size in EphA2-enriched and -depleted regions in CHO cells. The membrane protein LAT is known to be monomeric and serves as a control. Data for LAT has been previously published.^35^

To further investigate EphA2 interactions in EphA2-enriched and -depleted regions, we performed Förster Resonance Energy Transfer (FRET) experiments in HEK293T cells, using the Fully Quantified Spectral Imaging (FSI)-FRET method, with EphA2-mTurquoise and EphA2-eYFP as a FRET pair.^30^ The cells were activated with ephrinA1-Fc and imaged with a 2-photon microscope to acquire full fluorescence emission spectra with pixel-level resolution. A scan was performed at λ_1_ = 840 nm to excite the donor, mTurquoise, and a second scan was performed at λ_2_ = 960 nm to excite the acceptor, eYFP (**Figure 5 b and c**). The fluorescence emission spectra of pixels belonging to EphA2-enriched and -depleted regions are then spectrally unmixed to obtain 1) donor concentrations, 2) acceptor concentrations and 3) FRET efficiencies. **Figure 5 a-c** shows a typical cell and its associated data. The FRET binding curves, derived from the analysis of 200 EphA2-enriched and 213 EphA2-depleted regions are shown in **Figure 5 d and e**. The FRET efficiency increases as a function of acceptor concentrations as expected, as EphA2 is known to undergo concentration-dependent homoassociation in the cell membrane (**Figure 5 d**).^36^ The FRET binding curves overlap for raft and non-raft regions. The donor:acceptor expression ratios are shown in **Figure 5 e**. The ratio of EphA2 concentrations in EphA2-enriched to EphA2-depleted regions is 2.3 ± 0.03 (range: 0.5 – 5.0) (**Figure 2 a**).

**Figure 5.**
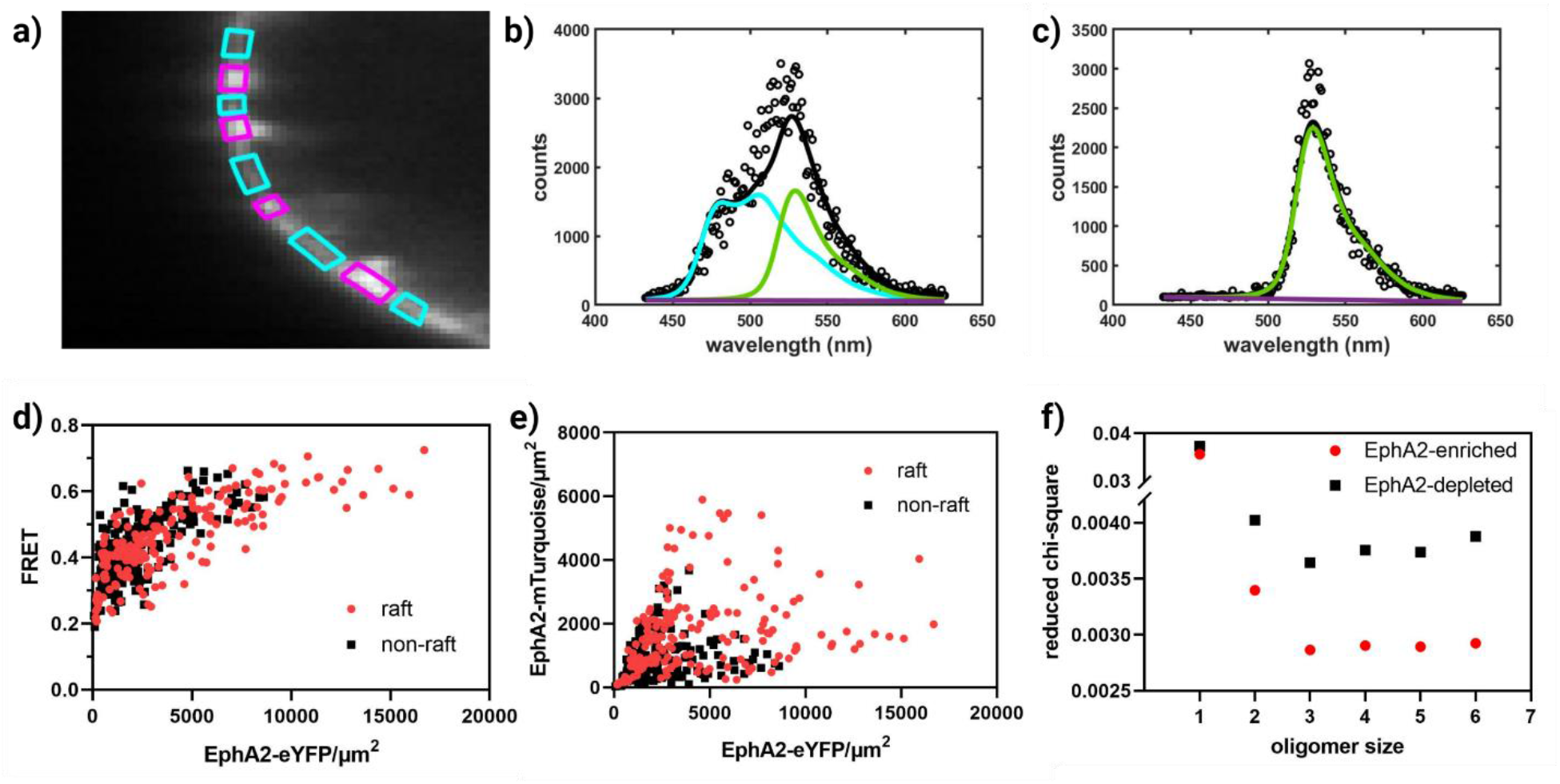
FRET data for EphA2-enriched and -depleted regions in HEK293T cells. a) – c) An illustration of the Fully Quantified Spectral Imaging (FSI) method.^30^ a) A HEK293T cell expressing EphA2-mTurquoise and EphA2-eYFP. Pixels within a selected membrane region of EphA2-enriched (purple polygon) and EphA2-depleted (turquoise polygon) regions are chosen for analysis. b), c) A single pixel’s fluorescence emission spectrum (black circles) is decomposed as a linear sum (black line) of donor (blue line), acceptor (green line), and background (purple) contributions in the FRET (b) and acceptor scans (c). d) FRET as a function of acceptor (EphA2-eYFP) concentration. Every data point corresponds to one region in a cell, as shown in a). e) Expression of the donor EphA2-mTurquoise and the acceptor EphA2-eYFP. f) The reduced chi-squared error versus oligomer order for monomer-only, monomer-dimer, monomer-trimer, monomer-tetramer, monomer-pentamer, and monomer-hexamer models. The model that best fits the data is the trimer model. d – f) Data for EphA2-enriched regions is shown in red and data for EphA2-depleted regions in black.

Following published protocols, we interpret the FRET data within the context of thermodynamic models based on the kinetic theory of FRET.^37–38^ Thermodynamic models are built for different types of oligomerization (n = 1-6) and the model that best describes the data is chosen to represent the data. The results of this MSE analysis are shown in **Figure 5 f**. We see that the MSE is minimized for n=3.

First, we note that the models used in the analysis assume a well-defined oligomer size, which appears to not be the case for EphA2, as suggested by the N&B data (**Figure 4**). Second, we note that this approach has been shown to reliably distinguish between dimers and higher order oligomers (n>2) but cannot reliably identify the exact oligomer size (n=3,4,5,…).^39^ Thus, the conclusion that we draw from the FSI FRET and N&B data is that the oligomer size of EphA2 exhibits a high degree of heterogeneity with a mean oligomer size of 3-4, both in EphA2-enriched and EphA2 depleted areas.

With this knowledge in hand, we fixed n=3 to construct oligomerization curves from the FRET data, in order to gain insights into the effects of the differential EphA2 concentration on EphA2 association (**Figure 6**). Notably, changes in the exact n does not change the oligomerization curve^39^, it only changes the fit parameters. The analysis and the underlying equations are briefly discussed in Materials and Methods and are described in detail in^30^. The oligomerization curves for EphA2 in EphA2-enriched and -depleted regions almost completely overlay and are well within the 68% confidence intervals of each other (dotted lines). The apparent association constant K_oligo,app_, i.e. the receptor concentration at 50% oligomeric fraction, can be read off the binding curves and is about 250 receptors/μm^2^. Such an oligomerization curve is useful if the expression of the receptor is known in biological context. There are reports of EphA2 expression of 600 receptors/μm^2^ in lung cancer lines ^40–41^; the curve in **Figure 6** therefore predicts that ~70% of the EphA2 will be oligomeric at this concentration.

**Figure 6.**
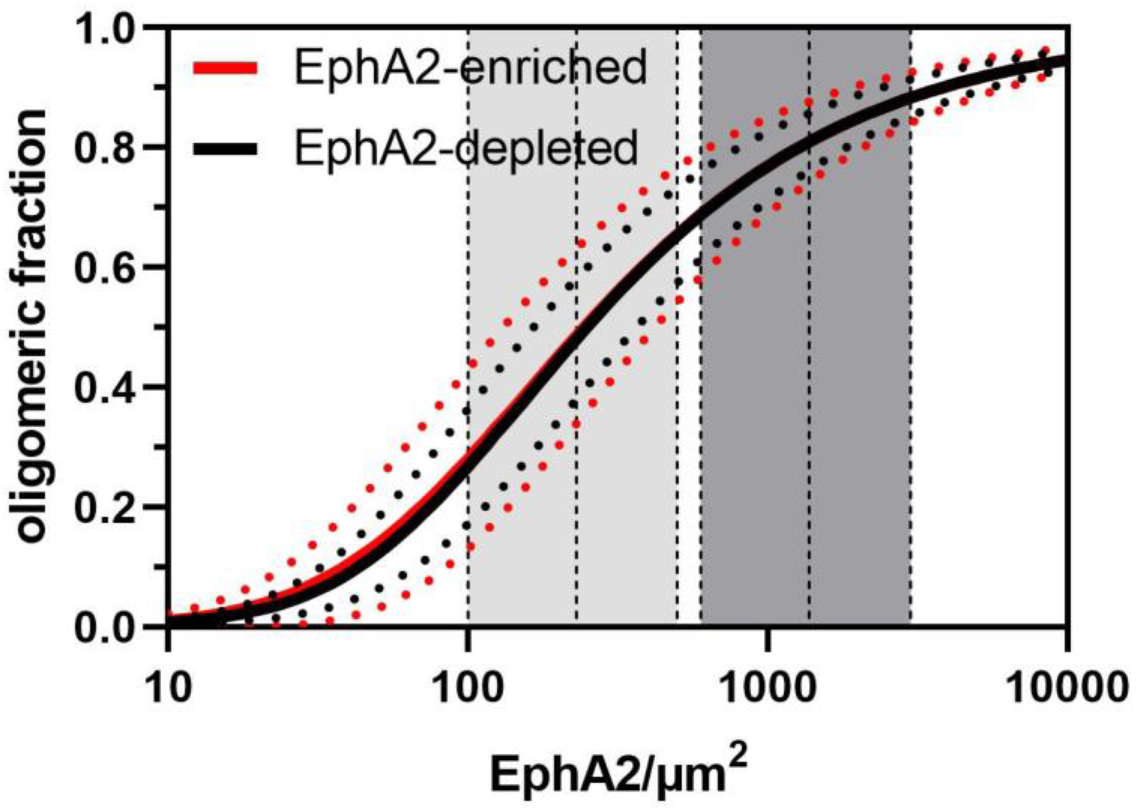
Oligomeric fraction as a function of EphA2 concentration for EphA2-enriched (red) and EphA2-depleted regions (black). The solid line is the best fit. The dotted lines around the fit show the 68% confidence interval of the fit. The vertical dashed lines indicate the EphA2 concentrations discussed in the text for cells with high EphA2 expression levels (dark grey shaded area) and cells with lower EphA2 expression levels (light grey shaded area).

The curve in **Figure 6** can tell us how the oligomeric fraction changes from EphA2-depleted to EphA2-enriched regions. On average, the EphA2 enrichment ratio is 2.3 ± 0.03 (range: 0.5 – 5.0) in the studied HEK293T cells (**Figure 2 a**). If we assume a concentration of 600 EphA2/ μm^2^ in the depleted regions, a 2.3 times enrichment in EphA2 concentration, from 600 to 1380 receptors/μm^2^ (**Figure 6**, from left border to middle dashed line of dark grey shaded area), leads to ~10% increase in the oligomeric fraction. In the extreme case of a 5 times EphA2 enrichment this would lead to a ~20% increase in the oligomeric fraction (right border of dark grey shaded area). However, EphA2 is known to be overexpressed in lung cancer, and thus normal EphA2 expressions will be lower. Assuming EphA2 receptor expression levels of 100 receptors/μm^2^ in the depleted domains (left border of light grey shaded area), we see that the average 2.3 times enrichment (middle dashed line, light grey shaded area) leads to a 22% increase in oligomeric fraction, as compared to EphA2-depleted domains. In the case of 5 times enrichment, 40% more receptors would be in an oligomeric state in the enriched regions as compared to the depleted regions (right border of light grey shaded area). Thus, while the difference of the concentration of oligomeric, signaling competent EphA2 receptors between enriched and depleted regions is modest in cells with overall high EphA2 expression, such as cancer cells, these differences are likely significant in cells with normal EphA2 expression levels.

Next, we sought to investigate if the separation of the membrane into EphA2-enriched and -depleted regions may affect downstream signaling. Once RTKs are phosphorylated, adaptor proteins bind the phosphorylated tyrosine residues of the RTK.^42^ Often, the adaptors associate with the plasma membrane since they are lipidated, and are believed to distribute themselves between raft and non-raft regions.^43^ One such adaptor is SRC, which contains a tyrosine kinase domain, regulatory SH2 and SH3 domains, and a myristoylated N-terminus.^44^ This adaptor is targeted to the membrane via the myristoylated N-terminus (**Figure S2**). The SH2 domain of SRC binds to juxtamembrane tyrosine phosphorylated motifs of EphA2.^44^ SRC can then, in turn, further activate EphA2 or transmit the signal downstream.^45–46^

HEK293T cells were co-transfected with EphA2-YFP and SRC-mTurquoise. The cells were stimulated with 50 nM EphrinA1-Fc and imaged in a two-photon microscope. Two scans were performed, following the FSI-FRET methodology, to yield FRET, EphA2 and SRC concentration maps for the plasma membrane.^30^ High FRET values that increase as a function of EphA2 concentration (**Figure 7**) confirm the specific EphA2 – SRC interactions in the plasma membrane in both EphA2 enriched and depleted domains. We fit the data in **Figure 7** to the exact solution of a single binding-site model (equation 6) to extract 1) the SRC – EphA2 dissociation constant K_D_ and 2) the structural parameter 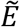, which depends on the average distance between the fluorescent proteins attached to the two interacting partners. **Figure 7** shows the best-fit curves for the EphA2-enriched (red line) and EphA2-depleted (black line) domains. The best-fit parameters are K_D,enriched_ = 2210 ± 780 proteins/μm^2^, 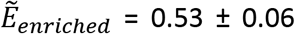 for the EphA2 enriched regions and K_D,depleted_ = 800 ± 390 proteins/μm^2^, 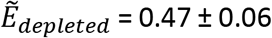 for the depleted regions, respectively. A two-tailed t-test of K_D_ demonstrates statistical significance of the dissociation constants calculated for the two types of domains (p < 0.0001). However, no significance was found for the values of the structural parameter 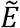 calculated for EphA2-enriched and -depleted domains (p = 0.46).

**Figure 7.**
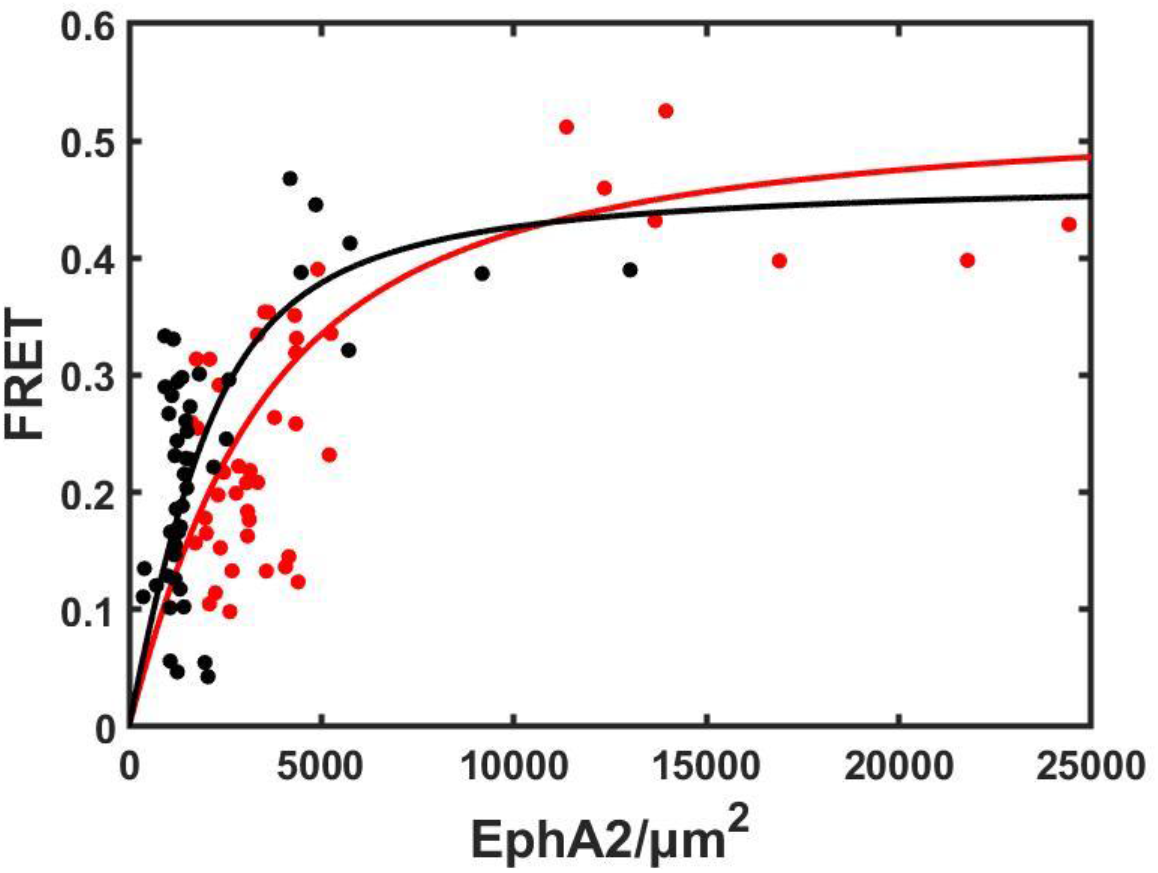
FRET reporting on the interactions between EphA2-eYFP and SRC-mTurquoise. FRET is shown as a function of EphA2 concentration for EphA2-enriched (red) and -depleted (black) regions. The high FRET values indicate direct interactions between EphA2 and SRC. The data for EphA2-enriched and -depleted regions were fit to the exact solution of a single binding-site model, given by equation 6. The two best-fit dissociation constants are different (p<0.0001), see text.

**Figure 8** further compares the spatial distribution of the enriched and depleted regions for EphA2 (**a**) and SRC (**b**) on the plasma membrane. The concentration ratio between enriched and depleted regions is 1.9 ± 0.1 (range: 0.8 – 3.7) for SRC and 2.2 ± 0.1 (range: 1.0 – 4.3) for EphA2 **(Figure 8 d**). The measured EphA2 ratio in this experiment (2.2 ± 0.1) is in agreement with the EphA2 ratio measured in the FRET experiment (2.3 ± 0.03). The enrichment of EphA2 is higher, as compared to SRC enrichment, and this effect is statistically significant (p < 0.0001, **Figure 8 d**). The PCC coefficient (equation 1) for EphA2 and SRC is 0.76 ± 0.05.

**Figure 8.**
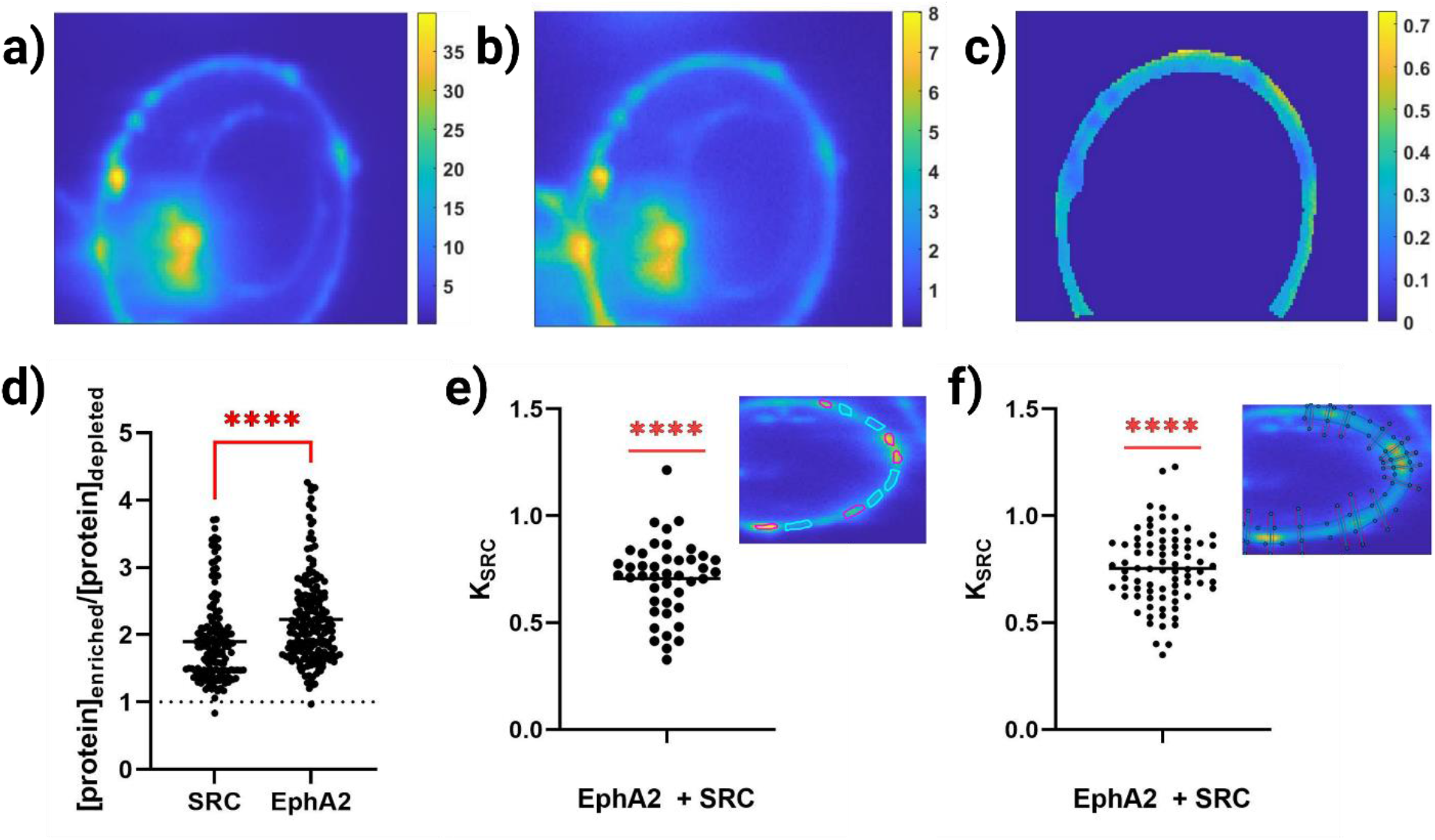
SRC abundance in EphA2-enriched and -depleted regions in a typical cell. a) Heatmap of EphA2 concentrations. b) Heatmap of SRC concentrations. c) Heatmap of the ratio of a) and b). d) Ratios of EphA2 and SRC concentrations in enriched and depleted regions. The degree of EphA2 enrichment is higher than SRC enrichment. e) and f) Comparison of SRC to EphA2 ratios in EphA2 enriched and depleted regions. K_SRC_ is given by equation (7). In e) calculations are performed using the ROI method (inset and see text). In f) calculations use the intensity profile method (inset and see text). Fewer SRC molecules are present per EphA2 in EphA2 enriched regions. This result can be confirmed visually by comparing a) and c): the locations of high EphA2 concentration in a) correspond to low SRC:EphA2 ratio in c).

To quantify whether there is a difference in adaptor protein partitioning to EphA2 enriched regions, we calculated K_SRC_, the ratio of SRC molecules per EphA2 receptor between EphA2-enriched and -depleted regions (equation (7)). To calculate K_SRC_, EphA2 and SRC concentrations were extracted from concentration maps with ROIs and intensity profiles as described above (**Figure 8 e and f**, inset). Both analysis methods, based on either ROIs (**Figure 8 e**, K_SRC_ = 0.71 ± 0.03) or intensity profiles (**Figure 8 f**, K_SRC_ = 0.75 ± 0.02), yield K_SRC_ values that are smaller than 1. Thus, fewer SRC molecules per EphA2 receptor are present in EphA2 enriched regions in agreement with the finding of reduced EphA2-SRC interactions in the EphA2 enriched regions. This result confirms what can be visually observed by comparing the EphA2 concentration map (**Figure 8 a**) to the SRC:EphA2 concentration ratio map (**Figure 8 c**): for each EphA2 enriched region (**Figure 8 a**), we see smaller SRC:EphA2 ratio as compared to the EphA2 depleted regions (**Figure 8 c**).

### Effects of TM sequence on EphA2 segregation

Studies in model systems have suggested that membrane protein incorporation into functional signaling domains is controlled by the sequence of the TM domain.^47^ Many of these prior studies have been performed with plasma membrane derived vesicles, produced via DTT and formaldehyde.^12, 48^ Here we investigated if the ratio of EphA2 concentrations in EphA2-enriched and -depleted regions in live cells is sensitive to the sequence of the EphA2 TM domain. We used two EphA2 mutants which have been previously shown to alter the phosphorylation of EphA2 kinase domain, and thus affect signaling.^49^ One is the triple mutant, G539I/A542I/G553I, referred to as the “HR mutant” in the literature.^49^ The other is a double mutant, G540I/G544I, referred to as the “GZ mutant” in the literature.^49^ We introduced these mutations into the EphA2 constructs labeled with fluorescent proteins at the C-terminus. Cells were transfected with these mutants, and their membranes were again observed to separate into EphA2-enriched and -depleted areas when ephrinA1-Fc was added. The ratios in concentrations between EphA2-enriched and -depleted areas are shown in **Figure 9**. We see statistically significant differences in the measured ratios (WT: 2.3 ± 0.03, range: 0.5 – 5.0; GZ: 1.6 ± 0.01, range: 0.3 – 3.1; and HR: 2.1 ± 0.02, range: 0.8 – 4.1) due to the sequence alterations. A multiple comparison ANOVA test revealed p-values < 0.0001 between all datasets.

**Figure 9.**
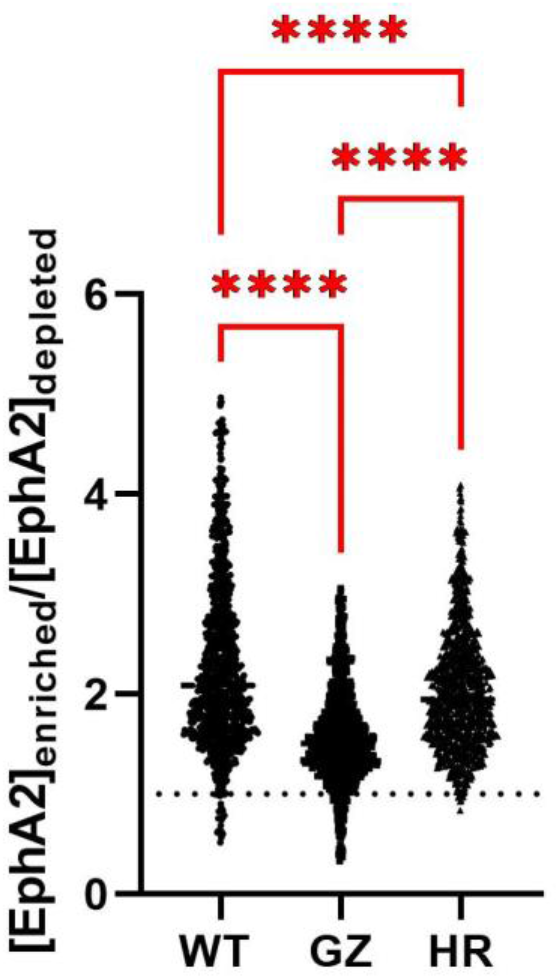
Effects of EphA2 TM sequence on EphA2 segregation. Concentration ratios for EphA2-enriched and -depleted regions for WT and the TM mutants GZ and HR. ANOVA shows highly statistical significance between the three datasets (p<0.0001).

## Discussion

Lipid rafts have been postulated to be highly ordered and tightly packed lipid microdomains that are enriched in gangliosides such as GM1.^19^ While micron-sized domains have been observed in model membranes and plasma membrane-derived vesicles using fluorescence microscopy, such structures have never been reported previously for the plasma membranes of mammalian cells. Instead, lipid rafts in plasma membranes have been proposed to be smaller than the diffraction limit of light and unstable^50–52^, and observable only if GM1 is cross-linked into large lattices using anti-cholera toxin antibodies. A recent report about the appearance of microscopic lipid domains in vacuole membranes of yeast in response to pH changes brought excitement and hope that lipid rafts can be observed in other biological membranes without lipid cross-linking treatments.^53^ Here we show for the first time that microscopic GM1-enriched domains can form in the plasma membrane of mammalian cells expressing the RTK EphA2. As GM1 is a known component of lipid rafts, our findings constitute a proof that micron-sized lipid domains can exist in the plasma membrane in biological context. Importantly, these GM1-enriched domains appear in response to a biological treatment, namely the stimulation with the ligand ephrinA1-Fc known to activate EphA2, rather than via GM1 cross-linking with anti-cholera toxin antibodies.

Lipid rafts have been proposed to form due to the phase behavior of lipids.^54–55^ Once the lipids are phase separated, specific proteins are believed to associate with the lipid rafts, while others associate with non-raft regions characterized by higher lipid disorder and mobility. Our results, however, suggest that the physical chemistry of the lipids is not sufficient for lipid raft formation because the observed GM1 separation occurs exclusively in response to ligand binding to a membrane receptor in the plasma membrane. Indeed, in the absence of ligand, we observe a homogeneous distribution of EphA2 and GM1 in the plasma membrane (**Figure 1**). Under these conditions, EphA2 is known to exist in a monomer-dimer equilibrium.^36^ Ligand binding to EphA2 leads to the formation of EphA2 oligomers that are larger than dimers, in addition to ligand-bound monomers and likely, ligand-bound dimers, which are all in dynamic equilibrium with each other.^36^ We can envision that EphA2 probably has preference for some lipids, and EphA2 oligomerization likely brings these lipids together. After this, we envision a nucleation of lipid domains that are enriched in GM1, which grow to reach stable sizes.

Proteins that play a role in signal transduction have been proposed to be enriched into the raft domains^3^, but such enrichment has never been directly quantified. Despite such lack of data, lipid rafts have been proposed to function as “signaling platforms”.^55^ Here we observe that ligand binding leads to the appearance of not only GM1-enriched domains, but also EphA2-enriched microdomains. This allowed us to quantify both EphA2 and GM1 colocalization and EphA2 enrichment. We find that there is no complete colocalization of EphA2-enriched and GM1-enriched domains (PCC = 0.34 ± 0.04 for HEK293T and 0.33 ± 0.02 for CHO cells). However, we do find that GM1 partitions preferentially into EphA2-enriched regions and vice versa (K_p,raft_ > 1 for CHO and HEK293T cells) (**Figure 3**). Yet, EphA2-enriched and GM1-enriched domains are distinct from each other.

When comparing EphA2-enriched and depleted regions we find that the EphA2 concentration is, on average, only a few fold higher in EphA2-enriched regions than in EphA2-depleted regions (**Figure 9**), an effect that is much more modest than expected based on the concept of a “signaling platform”^3^. Note that we perform measurements at very high ligand concentration (50 nM ephrinA1-Fc), and thus all EphA2 molecules are expected to be ligand-bound, no matter whether they localize to EphA2-enriched or depleted domains. The EphA2 oligomer sizes that we measure in the N&B experiments vary from 1 to 21, and are the same in EphA2 enriched and depleted domains. Association constants describing lateral interactions are also the same, in EphA2-enriched and depleted domains. The only identifiable difference is the larger fraction of EphA2 molecules that are associated into oligomers in the enriched regions since the EphA2 concentration there is higher. Consistent with this view, we see that the FRET efficiencies measured in the EphA2 enriched regions tend to reach somewhat higher values because the EphA2 concentrations are higher. Given the binding curves that we measure in the FRET experiments, this increase in oligomeric fraction can be either modest (up to 20%) if EphA2 is overexpressed (as in lung cancer cells, 600 rec/μm^2^), or significant (up to 40%) if EphA2 expression is lower. Overall, these differences are entirely due to differences in EphA2 concentrations, not in EphA2 lateral association in the plasma membrane in response to ligand.

What can induce EphA2 concentration heterogeneities in the plasma membrane? One possibility is that specific lipid-protein interactions play a role in the process. If this is so, then mutagenesis of EphA2 TM domains can affect the distribution of EphA2 in the plasma membrane. In support of this view, we find that the exact sequence of EphA2 TM domains affects the separation of EphA2 into enriched and depleted domains (**Figure 9**). The differences due to engineered TM domain mutations are small, but statistically significant. The two TM mutants of EphA2 that we investigate here have been previously shown to affect EphA2 function, but the effects were small. It is thus possible that the modest effects that we observe here underlie the previously reported small differences in EphA2 activity due to the TM domain mutations.^49^ Noteworthily, these mutations do not affect EphA2 lateral association, as assessed by FRET (data not shown). While many questions about the role of protein sequences in maintaining membrane heterogeneities still remain, we note that our experimental approach can be used for in-depth investigation of the effect of TM domain sequences in protein and lipid segregation.

Furthermore, we directly show for the first time that membrane heterogeneities influence receptor signaling by studying the behavior of the signaling adaptor SRC in live cells. EphA2 associates with SRC in both EphA2 enriched and depleted regions. SRC also forms enriched and depleted regions which are strongly colocalized with EphA2 (PCC = 0.76 ± 0.05). However, the heterogeneity in the SRC distribution (ratio between SRC-enriched and -depleted regions: 1.9 ± 0.1) is smaller than the heterogeneity of EphA2 (ratio between EphA2-enriched and -depleted regions: 2.2 ± 0.1). Analysis of the spatial heterogeneity distributions of EphA2 and SRC suggest that fewer SRC proteins are bound to EphA2 in EphA2 enriched regions when compared to EphA2 depleted regions (K_SRC_ ~ 0.7). To investigate this possibility, we directly measure the EphA2-SRC dissociation constants in EphA2 enriched and depleted domains, and we find that the association is indeed stronger in the depleted regions. The reason for this differential association is currently unknown, but it may reflect different dispositions of the myristoylated SRC in the EphA2 enriched and depleted domains. Importantly, our approach to create resolvable domains has now set the stage for biophysical measurements that can answer such mechanistic questions. Further, this study opens new avenues for investigations into a variety of signaling platforms, as we anticipate that different adaptors with different lipid modifications will behave differently.

To gain further insights into the nature of EphA2 heterogeneity in the plasma membrane, we compared the formation of EphA2-enriched and depleted domains in cells, swollen cells and vesicles. We observed EphA2 segregation in all cases. Note that the vesicles have no cytoskeleton, and that the cytoskeleton is largely disrupted in cells under reversible osmotic stress. Thus, our data suggest that the cytoskeleton does not play a crucial role in establishing EphA2 heterogeneity. In fact, the largest EphA2 heterogeneities were observed in vesicles which completely lack cytoskeleton. Indeed, when performing experiments with vesicles, produced from cells via osmotic vesiculation, we observed that the concentrations are up to 20 times higher in the EphA2 enriched areas as compared to EphA2 depleted regions. This is an order of magnitude larger segregation as compared to cells. This observation suggests that the plasma membrane components are indeed capable of significant phase separation, in accordance with the original raft hypothesis. However, such big differences are not observed in cells, suggesting that the cell actively reduces membrane heterogeneities.

While cells have distinct lipid compositions for the inner and outer leaflet of the plasma membrane, the plasma membrane-derived vesicles are believed to have scrambled lipid composition because the proteins that maintain the asymmetry of the plasma membrane are not functional.^56^ In support of this view, it has been shown that phosphatidylserine, mainly found in the inner leaflet of healthy cells, is exposed on the outer surface of vesicles.^57^ Perhaps, the native lipid composition of the leaflets in the living cell suppresses EphA2 heterogeneities.

Studies of EphA2 segregation in formaldehyde vesicles brought further surprises. While 100% of the imaged cells, swollen cells and osmotically-derived vesicles exhibited EphA2 enriched and depleted domains, about 70% of the formaldehyde vesicles did NOT exhibit any EphA2 segregation. For the subset of formaldehyde vesicles which phase separated, we recorded EphA2 intensity ratios for enriched and depleted domains that are smaller than the ratios measured for osmotically-derived vesicles. The reason for the observed differences between the two types of vesicles is unknown, but it may be due to formaldehyde-induced cross-linking of components in the formaldehyde vesicles. Yet, curiously, it is the lipids in the formaldehyde vesicles that are known to readily phase separate into domains under some conditions.^58^

In summary, this work documents new observations and new insights into plasma membrane heterogeneities and their role in cell physiology. First, the experiments described here outline a simple approach to induce both EphA2 and GM1 enriched and depleted domains. The latter are distinct and readily observable in a fluorescence microscope, and offer a tool to investigate the behavior of molecules believed to partition differentially between lipid raft and non-raft domains. Second, the data presented here amend the notion of the raft as a “signaling platform”^3^. We do not observe complete colocalization between the signaling receptor EphA2 and the lipid raft constituent GM1. Thus, the EphA2 domains themselves likely serve as signaling platforms. By measuring dissociation constants, we demonstrate that EphA2 oligomerization is the same in EphA2-enriched and -depleted domains. However, EphA2 interacts preferentially with its downstream effector SRC in EphA2-depleted domains. Further studies with other signaling molecules are needed for comprehensive understanding of signaling platforms organization.

The observation of microscopic GM1-enriched domains in live cells in response to a biological ligand gives us unprecedented opportunities to study the biology of rafts. We hope that the new knowledge that we will gain through the use of this platform will lead to new ways to manipulate processes in biological membranes, and perhaps to a new generation of smart drugs that can finetune cell signaling by modulating membrane heterogeneities.

## Materials and Methods

### Plasmid constructs

A plasmid encoding for the human receptor EphA2 tagged with a fluorescent protein (eYFP or mTurquoise) separated by a flexible 15 amino acid (GGS)5 linker in pcDNA3.1(+) was created as described ^36^. The glycine zipper (GZ: G540I, G544I) and heptad repeat (HR: G539I, A542I, G553I) variants were created using the QuikChange II Site-Directed Mutagenesis Kit according to manufacturer’s instructions (Agilent Technologies, #200523). All plasmids were sequenced to confirm their identity (Genewiz).

The construct for the adaptor protein mTurquoise-SRC-N-18 (SRC with mTurquoise fused to its C-terminus via a linker) was a gift from Michael Davidson (Addgene plasmid # 55560; http://n2t.net/addgene:55560; RRID:Addgene_55560).

The constructs were amplified in competent DH5α *E. coli* cells and purified using Qiagen’s HiSpeed Plasmid Midi Kit (# 12643).

### Cell culture and transfection

HEK293T cells (ATCC) were cultured in Dulbecco’s modified eagle medium (Gibco, #31600034) supplemented with 10% fetal bovine serum (HyClone, #SH30070.03), 20 mM D-Glucose and 18 mM sodium bicarbonate at 37 °C in a 5% CO_2_ environment. Chinese hamster ovary (CHO) cells were cultured in Dulbecco’s modified Eagle medium (Gibco, #31600034) supplemented with 10% fetal bovine serum (HyClone, #SH30070.03), 1 mM nonessential amino acids, 10 mM D-glucose, and 18 mM sodium bicarbonate at 37 °C in a 5% CO_2_ environment.

For cellular assays, the cells were seeded in 35 mm glass coverslip, collagen-coated imaging dishes (MatTek, P35GCOL-1.5-14-C) at a density of 2.5*10^5^ (HEK293T) or 1*10^5^ (CHO) cells per dish. 24 hours after seeding, Lipofectamine 3000 (Invitrogen, #L3000008) was used for transient transfection according to the manufacturer’s protocol. For domain characterization, colocalization, and N&B experiments, the cells were transfected with 1 – 2 μg EphA2-mTurquoise plasmid DNA. For FRET association experiments, cotransfections were performed with 2 – 3 μg total plasmid DNA in a 1:3 donor (EphA2-mTurquoise):acceptor (EphA2-eYFP) ratio. For adaptor protein recruitment experiments, the cells were cotransfected with 1 – 2 μg EphA2-eYFP and 1 – 2 μg mTurquoise-SRC. Single transfections were performed for the spectral unmixing of cotransfected cell images using 1 μg of plasmid DNA. 12 hours after transfection the cells were washed twice with phenol-red free, serum-free starvation media and then serum starved for at least 12 hours.

For vesicle experiments CHO cells were seeded into a 6-well plate at a density of 2*10^4^ cells per well. 24 hours after seeding, the cells were transfected with 1 – 2 μg EphA2-mTurquoise plasmid DNA using FuGene HD (Promega, #E2311) according to the manufacturer’s protocol. For osmotically-derived salt vesicles, the cells were rinsed 36 h after transfection with 1 ml of 30% PBS and incubated at room temperature for 1 minute. The rinse was performed two times. Then 1 ml of osmotic vesiculation buffer (**Table 1**) was added to each well and the plate was incubated for 13 h at 37 °C in a 5% CO_2_ environment. After incubation, the vesicles were carefully transferred to an 8-well glass bottom chamber slide (ibidi, #80827) for imaging. For formaldehyde/DTT vesicles, the cells were washed twice with 1xPBS 48 h after transfection. Then, 1ml of freshly prepared formaldehyde/DTT buffer (**Table 1**) was added and the cells were incubated for 1 h at 37 °C in a 5% CO_2_ environment. After incubation, the PFA vesicles were carefully transferred to an 8-well glass bottom chamber slide (ibidi, #80827) for microscope analysis.

**Table 1.** Vesicle buffer recipes.

| Buffer | Osmotic buffer | Formaldehyde/DTT buffer |
| --- | --- | --- |
|  | 200 mM NaCl | 0.5 mM MgCl<sub>2</sub> |
| **Ingredients** | 5 mM KCl | 0.75 mM CaCl<sub>2</sub> |
|  | 0.5 mM MgCl<sub>2</sub> | 25 mM formaldehyde |
|  | 0.75 mM CaCl<sub>2</sub> | 0.5 mM DTT |
|  | 100 mM Bicine | PBS (1.06 mM KH<sub>2</sub>PO<sub>4</sub> + 2.97 mM Na<sub>2</sub>HPO<sub>4</sub> + 155 mM NaCl) |
| **pH** | 8.5 | 7.4 |

### Microscope imaging

For domain characterization and colocalization experiments, the starvation media was replaced with 1 ml fresh starvation media, supplemented with 0.3 μg Cholera Toxin Subunit B labeled with Alexa Fluor 555 (V-34404, Vybrant) and 50 nM dimeric EphrinA1-Fc (R&D Systems, #6417-A1). Control experiments were performed without ephrinA1-Fc. The cells were imaged immediately with a TCS SP8 confocal microscope (Leica Biosystems, Wetzlar, Germany) equipped with a HyD hybrid detector and a diode laser. Two scans were acquired, an ‘EphA2’-scan (λ=448nm, emission window: 460 – 520nm), where mTurquoise bound to EphA2 is excited, and a ‘GM1’-scan (λ=552nm, emission window: 560 – 610nm), where Alexa Fluor 555 bound to CTB is excited. Excitation wavelengths and emission windows were chosen so that spectral bleedthrough is absent in the experiment. Images were acquired at 1.5% laser power with a 100Hz scanning speed. Each imaging session did not exceed one hour.

FRET association and adaptor protein recruitment experiments were performed according to the FSI-FRET methodology.^30^ Before the experiment, the starvation media was replaced with a pre-warmed (37 °C) 1:9 serum-free media:diH_2_O, 25 mM HEPES solution. This reversible osmotic swelling unwrinkled the cell membrane such that it 1) removed membrane indentations and ruffles that could be misidentified as rafts, 2) converted the membrane topology into a sphere, allowing for precise calculation of protein concentrations, and 3) prevented EphA2 ligand-induced endocytosis.^30^ The cells were allowed to stabilize for 10 min at room temperature prior to imaging. FSI-FRET imaging was performed with a two-photon microscope equipped with the OptiMis True Line Spectral Imaging system (Aurora Spectral Technologies, WI).^37, 59^ Two microscope scans per cell were acquired – an acceptor excitation scan (to measure acceptor fluorescence, λ=960 nm) and a donor excitation scan (to measure both donor and sensitized acceptor fluorescence, λ=840 nm). A scan is composed of 300×440 pixels, and each pixel contains a full fluorescence spectrum in the range of 420 – 620 nm. Each imaging session did not exceed 1 hour. The measured pixel-level intensities of the images were converted to concentrations with calibration solutions of purified fluorescent protein as previously described^30^. The soluble monomeric eYFP and mTurquoise calibration solutions were produced following a published protocol ^60^.

Number and Brightness (N&B) data were acquired and analyzed as described previously ^35^. N&B measures the ratio of the variance due to the fluorescence intensity fluctuations over time to the average intensity, which is different for different types of oligomers. ^34^ Experimentally, the fluorescence intensity fluctuations are measured by rapidly taking an image stack of the same region of a cell, and then computing the mean fluorescence intensity and the variance across the stack for each pixel. This allows for the calculation of the apparent molecular brightness, ε_app_. The brightness does not depend on the concentration and scales linearly with the oligomer size. Thus, the oligomer size of EphA2 can be calculated by normalizing the molecular brightness measured for EphA2 to the molecular brightness of LAT, a monomer control.

N&B imaging was performed on a TCS SP8 confocal microscope (Leica Biosystems, Wetzlar, Germany) equipped with a HyD hybrid detector in photon counting mode. The cells were excited with a 448 nm diode laser at 0.1% of the maximal power, and the emission spectrum windows was the default CFP setting. An image stack of 150 images was taken over approximately 90 seconds, with each image being 256×256 pixels. The pixel dwell time was 5.4 μs. A 18x optical zoom was used, corresponding to a 40.2 nm pixel length. Fluorescence intensities were converted into concentrations by creating calibration curves by measuring purified mTurquoise standards of known concentrations under the same imaging conditions, as described in detail previously^30^.

### Image analysis

For intensity ratio analysis, EphA2/GM1-enriched and -depleted regions were manually outlined by the researcher in the ‘EphA2’- or ‘GM1’-scan, respectively. The mean intensity values within these regions were used to calculate the intensity ratio between EphA2/GM1-enriched and -depleted domains. Ratios of intensity were always calculated from regions of the same cell. In the case of multiple enriched and depleted regions in a cell, each enriched region intensity was divided by all depleted region intensities.

For colocalization analysis, the cell membrane was outlined in the ‘EphA2’-scan. Using the membrane ‘EphA2’- and ‘GM1’-scan pixel intensity values EphA2i and GM1i the Pearson correlation coefficient (PCC) is calculated:

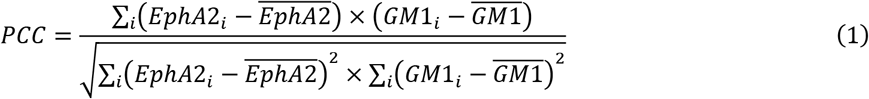

where 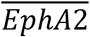 and 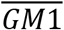 refer to the mean intensities of the ‘EphA2’-scan and the ‘GM1’-scan membrane regions, respectively.

To analyze the images of GM1 partitioning experiments, intensities of CTB-Alexa Fluor 555 in EphA2-enriched and EphA2-depleted regions were compared. Regions of interest were identified in the ‘EphA2’-scan and manually outlined by the researcher (**Figure 3 a, ii**). The pixel-level ‘GM1’-scan intensities of the chosen regions were averaged and used to calculate the GM1 partitioning coefficient K_p,GM1_:

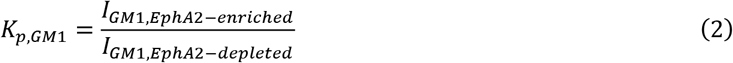

where I_GM1, EphA2-enriched_ is the CTB-Alexa Fluor 555 intensity inside the EphA2-enriched region and I_GM1, EphA2-depleted_ the CTB-Alexa Fluor 555 intensity inside the EphA2-depleted region right next to it.

To confirm the results of this analysis, a second analysis method was used. Here, lines are drawn across EphA2-enriched or -depleted regions (**Figure 3 b, i**). For each line across an EphA2-enriched region, two adjacent lines across EphA2-depleted regions were selected. The selected lines are then expanded by two parallel lines (red) and the average intensity profile of the expanded selection is computed inside the outlined area of interest (blue polygon) as seen in **Figure 3 b, i and ii**. The maximum intensities along the intensity profiles are chosen as I_GM1, EphA2-enriched_ or I_GM1, EphA2-depleted_. Equation 2 is then used to calculate K_p,GM1_. For control images without added ephrinA1-Fc, the membrane was homogenous and regions were randomly assigned as enriched and depleted. The image analysis for GM1 partitioning experiments was performed with custom-written Matlab code.

For EphA2 association experiments, the collected images were analyzed with the FSI-FRET software^30, 61^. In short, the fluorescence emission spectra for every pixel of the image were unmixed to yield donor (mTurquoise) and acceptor (eYFP) fluorescence contributions. The unmixed donor and acceptor contributions were then background corrected and integrated to give intensity values per pixel. These intensity values were then compared to calibration solutions of purified fluorescent protein to yield donor, acceptor concentrations, and FRET as previously described ^30, 60^. These concentration values were used to calculate concentration ratios for EphA2-enriched and -depleted regions for WT EphA2 and the TM mutants GZ and HR.

To analyze EphA2 homoassociation experiments, concentrations (EphA2-mTurquoise, donor and EphA2-eYFP, acceptor) and FRET values were calculated for EphA2-enriched and -depleted regions to yield FRET binding curves. The FRET occurring due to specific association of donors and acceptors, E_oligo_, can be described by Raicu’s kinetic theory of FRET ^38^:

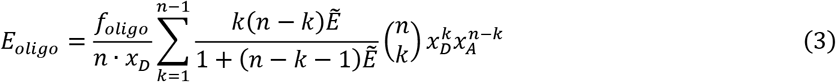

where f_oligo_ is the fraction of proteins in the oligomeric state, n the oligomer order (dimer, trimer, …), and 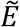 a structural parameter, which depends on the separation and orientation of the fluorophores. x_D_ and x_A_ are the donor fraction and acceptor fraction, respectively. foligo depends on the association constant K_a_ and the total receptor concentration [T] according to:

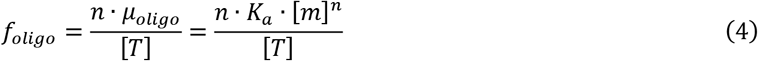

where [m] is the concentration of monomers.

However, the measured FRET efficiency has two contributions: 1) E_oligo_, FRET due to specific association of the EphA2 receptor (see above) and 2) stochastic FRET (E_stochastic_). Stochastic FRET, or proximity FRET, occurs when donor and acceptor approach each other by chance within close proximity (<100 Å) and is especially pronounced in the confined plasma membrane. To account for stochastic FRET, we add a term in equation 3 that reflects FRET due to random proximity of donors and acceptors ^30, 39, 62^:

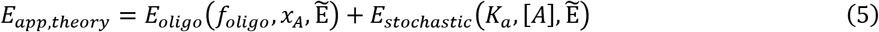

Stochastic FRET depends on the unknown parameters K_a_, 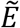, and the measured acceptor concentrations. A stochastic FRET library for n=2-6 has been published in previous work^39^ and was used to calculate the theoretical FRET efficiency, E_app,theory_, for the adjustable parameters n (2-6), K_a_, and 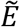. The theoretical FRET efficiencies for different n, K_a_, and 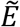 were then compared to the experimental measured ones, and the mean squared errors (MSEs) were calculated. The model with the lowest MSE was chosen to best represent the data and used to yield oligomeric fraction vs concentration plots.

For the analysis of binding between the adaptor protein SRC-mTurquoise and EphA2-YFP, the concentrations of SRC-mTurquoise, the concentration of EphA2-YFP, and the FRET efficiency for EphA2-enriched and -depleted regions were obtained using the FSI-software.^30, 61^ The resulting three dimensional datasets were fit with the exact solution of a FRET binding curve for a single site model:

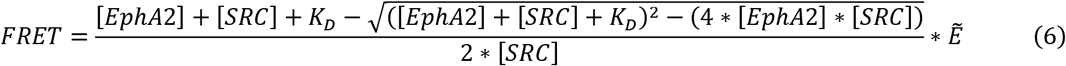

to yield K_D_, the EphA2 – SRC dissociation constant, and 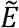, a structural parameter, which depends on the separation and orientation of the two fluorophores in the complex. A two-tailed t-test was performed to compare the best-fit K_D_ and 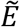 values of EphA2-enriched and -depleted regions. The significance cut-off was set to α=0.05. To plot the best-fit curve, a slice of the best-fit surface to the three-dimensional datasets at [SRC] = 2000 SRC/μm^2^ was extracted.

Furthermore, SRC:EphA2 concentration ratios were computed by selecting EphA2-enriched and -depleted regions on the EphA2 concentration map and used to calculate K_SRC_ according to:

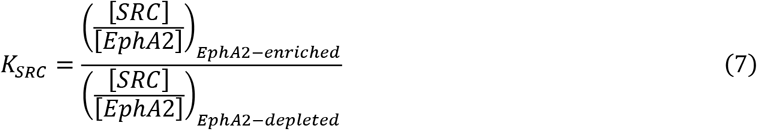

where [SRC] and [EphA2] are the measured SRC and EphA2 concentrations in EphA2-enriched or -depleted regions, respectively. Similarly to the colocalization analysis, the images were analyzed by 1) defining regions of interest with polygons and 2) selecting lines over EphA2-enriched and -depleted regions.

## Data Availability

All data are included in the manuscript

## Acknowledgments

The authors thank Dr. Elmer Zapata-Mercado for the actin images and Randall Rainwater for help with N&B experiments.

## Funding

Supported by NIH grants R01GM131374 (EBP and KH), R01GM068619 (KH), and NSF MCB 2106031 (KH)

## Conflict of interest

The authors declare no conflict of interest.

## Notes

### Competing Interest Statement

The authors have declared no competing interest.

