## Supplementary Figures for "Direct quantification of ligand-induced lipid and protein microdomains with distinctive signaling properties"

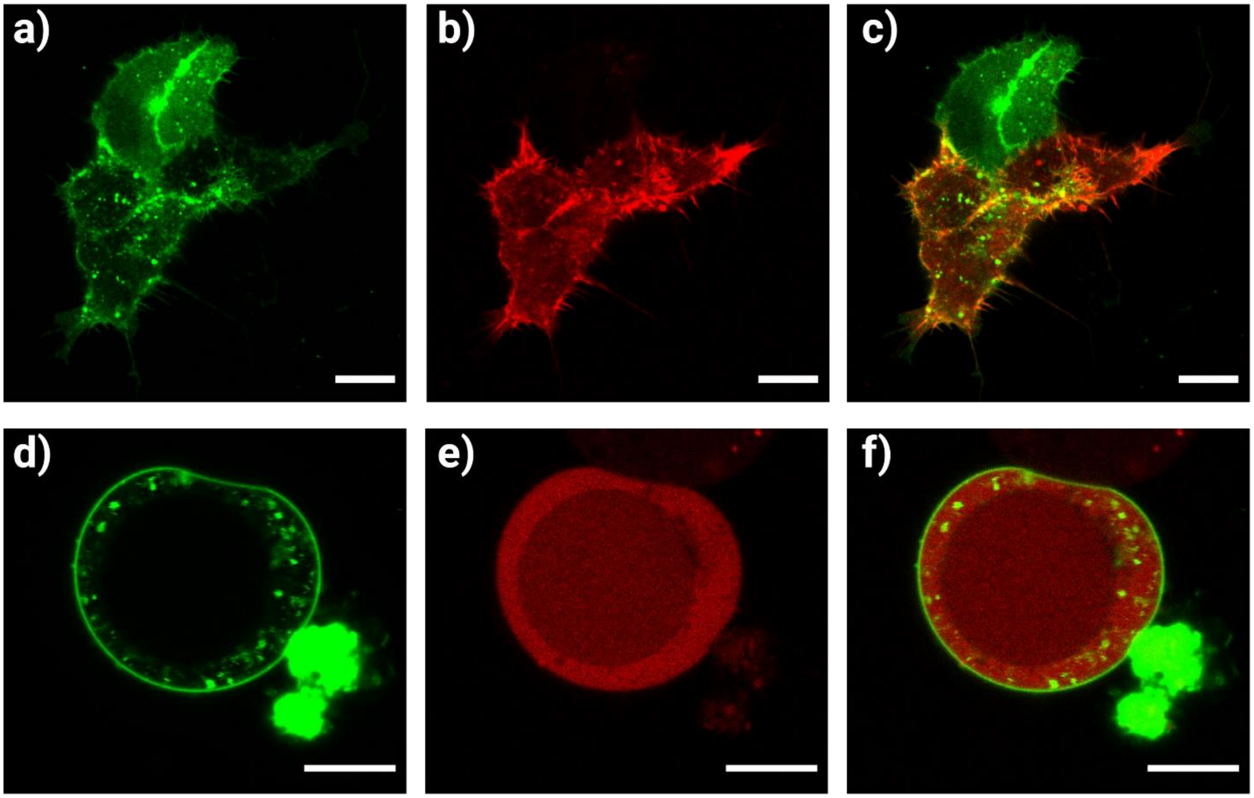


**Figure S1**. Cortical actin is disrupted in cells under osmotic stress. Cells expressing EphA2-eYFP (green, a and d) and Lifeact-mCherry, a marker to visualize actin (red, b and e), are shown, as well as the overlay of the two channels (c and f). Top row: In normal media, the intact actin network can be seen in cells (b). Bottom row: Cells under osmotic stress. Only homogenous fluorescence can be observed in (e), suggesting that the actin network in these cells is depolymerized. The scale bar represents 10 µm.

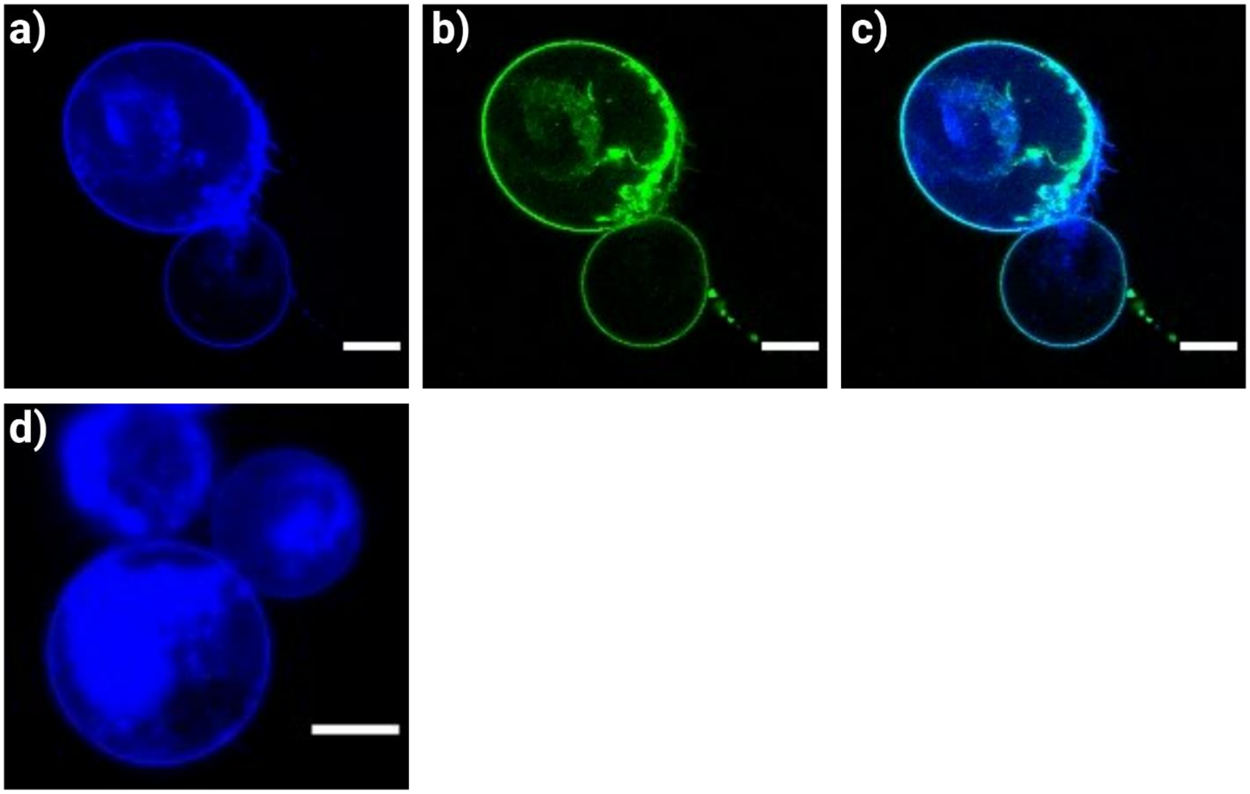


**Figure S2**. SRC recruitment to the cell membrane in the absence of ligand. Top row: Swollen HEK293T cells expressing SRC-mTurquoise (a) and EphA2-eYFP (b). The overlay of the two channels is shown in c. Even in the absence of ligand, the adaptor protein is recruited to the plasma membrane. Second row: Swollen HEK293T cells expressing SRC-mTurquoise (d). Even in the absence of EphA2, SRC is localized at the plasma membrane, likely because of its myrostoylated N-terminus. The scale bar represents 10 μm.
